# Triclabendazole and sutezolid are not effective *in vivo* against *Plasmodium berghei*

**DOI:** 10.64898/2026.08.05.743170

**Authors:** Jérôme Dormoi, Rémy Amalvict, Leo Millot, Bruno Pradines

**Author notes:** Correspondance: Jérôme Dormoi.

## Abstract

Drug repositioning has emerged as an attractive strategy to accelerate the development of new antimalarial therapies, particularly by evaluating compounds already used against pathologies co-endemic with malaria. This approach offers the advantage of leveraging existing pharmacokinetic, toxicological, and safety data, thereby potentially shortening the drug development pipeline. However, transposing a compound from its original therapeutic indication to an antimalarial use is far from straightforward: differences in target biology, parasite stage specificity, pharmacodynamic requirements, and host-parasite interactions can result in a loss of efficacy despite promising in vitro or structural rationale. Rigorous in vivo validation therefore remains indispensable before any repositioning hypothesis can be considered translationally relevant.

In this context, we evaluated the blood-stage antimalarial activity of triclabendazole, an antihelminthic drug used against co-endemic fascioliasis, together with its metabolite triclabendazole sulfoxide, and sutezolide, an oxazolidinone antibiotic, in a murine model of *Plasmodium berghei ANKA* infection following oral administration. None of the three compounds demonstrated significant antimalarial activity under these experimental conditions, contradicting a previously published repositioning hypothesis.

Beyond these specific findings, our study is deliberately framed within the 3Rs principles (Replacement, Reduction, Refinement) governing animal experimentation. We argue that publishing negative *in vivo* results is not only scientifically legitimate but ethically necessary: sharing such data allows research teams working on similar preclinical models to build on existing knowledge, avoid unnecessary experimental duplication, and ultimately reduce the number of animal procedures performed across the field. We advocate for wider dissemination of negative outcomes in antimalarial drug repositioning research as a concrete contribution to more responsible and efficient use of animal models in preclinical pharmacology.

Graphical Abstract

## Introduction

Malaria remains one of the world’s most devastating parasitic diseases. It is caused by protozoan parasites of the genus *Plasmodium* and is transmitted to humans through the bite of infected *Anopheles* mosquitoes. Despite substantial progress in malaria control over the past two decades, the disease continues to represent a major global public health challenge. In 2024, an estimated 282 million malaria cases and approximately 610,000 malaria-related deaths were reported worldwide. The burden remains overwhelmingly concentrated in sub-Saharan Africa, where the World Health Organization (WHO) African Region accounted for approximately 95% of all cases and 96% of malaria deaths. Children younger than five years remain the most vulnerable population, representing nearly 80% of malaria-related fatalities [1–3]. Beyond its mortality, malaria is also responsible for considerable morbidity, impaired childhood development, reduced productivity, and substantial economic costs for endemic countries and their healthcare systems [4, 5].

Achieving malaria elimination requires overcoming several interconnected challenges. Among the most important are the effects of climate change on vector distribution, the increasing resistance of mosquito populations to insecticides, behavioral adaptations of malaria vectors following the widespread deployment of long-lasting insecticidal nets (LLINs), the limited sustainability of current preventive strategies, the emergence and spread of resistance to antimalarial drugs, the limited availability of highly effective vaccines, insufficient community health education, and the socioeconomic conditions that sustain malaria transmission [6–10]. Among these challenges, the continuous emergence of drug-resistant *Plasmodium falciparum* strains represents one of the greatest threats to malaria control and highlights the urgent need to identify novel antimalarial compounds with new mechanisms of action.

The discovery and development of new antimalarial drugs is, however, a lengthy, expensive, and high-risk process. Drug repurposing has therefore emerged as an attractive alternative strategy to accelerate antimalarial drug discovery. This approach relies on the evaluation of approved or clinically characterized compounds for new therapeutic indications. Because these molecules have already undergone extensive pharmacological, pharmacokinetic, and toxicological characterization, drug repurpose can substantially reduce both development time and costs while increasing the likelihood of successful clinical translation. More recently, advances in computational biology have further strengthened this strategy through the integration of artificial intelligence, machine learning, molecular docking, structural biology, and large-scale in silico screening approaches. These technologies enable the identification of previously unrecognized interactions between existing compounds and parasite targets, thereby facilitating the prioritization of candidates for experimental evaluation.

Among potential repurposed compounds, sutezolide (STZ), triclabendazole (TBZ) and one of its main derivatives, attracted our attention following the review by Fontinha *et al.* (2020), which suggested evaluating this benzimidazole derivative against the liver stage of *P. falciparum* infection rather than against the erythrocytic stage (11). To our knowledge, however, no experimental study has investigated the antiplasmodial activity of TBZ or its major metabolites against the blood stages of *P. falciparum*. This absence of experimental evidence is particularly noteworthy because TBZ belongs to the benzimidazole family, a chemical class that has already demonstrated antiplasmodial properties. Several benzimidazole derivatives have shown activity against different developmental stages of *Plasmodium*, mainly through interference with microtubule assembly and parasite cell division [11]. Furthermore, recent studies continue to report the development of novel benzimidazole-based compounds with promising antiplasmodial activity and improved structure-activity relationships [12, 13]. Collectively, these observations support further investigation of TBZ as a potential antimalarial candidate. TBZ is currently the treatment of choice for human fascioliasis and remains the only drug recommended by the WHO against both immature and adult stages of *Fasciola hepatica*.

Following oral administration, TBZ undergoes extensive hepatic metabolism, leading primarily to the formation of two active metabolites through sequential sulfoxidation and sulfonation: triclabendazole sulfoxide (TBZOX) and triclabendazole sulfone (TBZONE). These metabolites account for most of the circulating drug exposure in vivo and contribute substantially to the therapeutic efficacy observed against liver flukes. Consequently, evaluating TBZ alone would provide only a partial assessment of its potential antiplasmodial activity, making it essential to investigate its major metabolites alongside the parent compound. In flukes, it has been proposed that TBZOX act by inhibiting microtubule polymerization [14]

In another hand, STZ is an experimental antibiotic of the oxazolidinone class proposed to fight against multidrug-resistant and extensively drug-resistant tuberculosis [15]. Its action against bacteria, by inhibiting bacterial messenger RNA translation by binding to the 50S ribosomal subunit, has aroused our scientific curiosity. Like TBZ, the active form of STD appears after sulfoxidation to produce sutezolide sulfoxide. Given that doxycycline was previously and effectively repurposed for malaria therapy [16], we focus on a antibacterial drug which also act on protein synthesis even if doxycycline act differently by inhibiting bacterial messenger RNA translation by binding to the 50S ribosomal subunit and by blocking protein translation in the essential apicoplast organelle in *Plasmodium* [16]

In the present study, we evaluated the *in vitro* antiplasmodial activity of STZ, TBZ and one of its main metabolites, TBZOX, against the asexual blood stages of *P. falciparum*. Contrary to expectations based on the pharmacological profile of benzimidazole derivatives and oxazolidinone, none of the three compounds demonstrated significant antiplasmodial activity under our experimental conditions. These negative findings provide important evidence regarding the limitations of triclabendazole repurposing for malaria treatment and contribute to refining future drug-repositioning strategies. We further discuss the implications of these results considering the current knowledge on benzimidazole derivatives and their proposed mechanisms of action against *Plasmodium* parasites.

## Material and methods

### Solvent

Due to the limited solubility of the powders in aqueous media [17–19], all compounds were dissolved in pharmaceutical-grade oil and subjected to sonication under sterile conditions. The pharmaceutical-grade oil was kindly supplied by the Institut de Chimie Radicalaire – Nicolas Primas (PharmD, PhD). Sonication was performed using a Vibra-Cell™ 75185, Bioblock Scientific® at 50% amplitude 50 KHz, maintained in water with crushed ice, for 2 minutes).

### Drugs and Reagents

Artesunate (ART; MERCK, A7986-10MG) was employed as a positive control. The investigational drugs, TBZ (Santa Cruz Biotechnology, sc-213105), TBZOX (Santa Cruz Biotechnology, sc-475819), and STD (MyBioSource/Clinisciences, MBS387349-100mg), were prepared and stored according to manufacturer instructions. All drugs were administered orally using an adaptive device (Instech Plastic Feeding Tubes, 20 x 38 mm) to minimize injury during gavage. The dosing regimens were as follows: ART at 3 mg/kg daily for three days; STD at 50 mg/kg daily for six days [20, 21]; TBZ and TBZOX at 50 mg/kg daily for five days, with the latter two drugs given in three doses every four hours, totaling 50 mg/day [22, 23]. The pharmaceutical-grade oil was used as negative control.

### Parasite Strain

Antimalarial efficacy was assessed using the chloroquine-sensitive *Plasmodium berghei* ANKA (*Pb*A), strain MRA-671 obtained from the BEI Resources (NIH-supported program managed by ATCC). The parasite stock was maintained via serial passage through mice.

### Animal Model

Mice were housed in a 1290D box (Technoplast, Lyon, France) respecting a minimal volume for each mouse (840 cm^3^). Male and female BALB/c albino mice, weighing 20–22 g (aged 7–8 weeks old) from the Charles Rivers Laboratory (Saint Germain Nuelles, France) were housed under standard and controlled constant laboratory conditions (19–22 °C, relative humidity around 60%), and were fed a normal diet (FT-SAFE-A04, Safe, Augy, France) with water ad libitum. Mice were randomly allocated into groups to minimize bias. Mice were monitored for general health throughout infections to ensure they did not reach IACUC endpoints. Shelters and elements to gnaw were provided (JBM022, JBS021, JCEHORA500, Serlab, Montataire, France), and tube handling (JMTT3322, Serlab) was used to manipulate mice and change mice from one box to another following the National Centre for the Replacement, Refinement and Reduction of Animals in Research (NR3C) recommendations, “How to pick up a mouse”. Heats pads to increase temperature, were supplied as soon as mice exposed immobilization phase, wet diets and long baby bottle teats were used from D2 to minimize effort after infection.

### Inoculum Preparation

A donor mouse was inoculated intraperitoneally with *P. berghei* ANKA (*Pb*A) parasites. Parasitemia was monitored daily, and blood smears were prepared on day five post-infection. Mice exhibiting parasitemia levels between 10–15% were selected for inoculation. Under anesthesia (ketamine 100 mg/kg; xylazine 10 mg/kg), 0.5 mL of blood was collected via the retro-orbital sinus using a heparinized syringe, transferred into a sterile tube, and diluted to 15 mL with 0.9% NaCl solution. Each recipient mouse was then inoculated intraperitoneally with 0.2 mL of the blood suspension, containing approximately 10^7 infected erythrocytes.

### Antimalarial Treatment Protocol

In contrast to the classical Peters’ 4-day suppressive test [24], this study employed a curative approach, initiating treatment upon detection of parasitemia ≥1% via blood smear analysis, typically 2 days post-infection, consistent with previous publications [25–27].

### Parasitemia count

To calculate the percentage of parasitemia on a thin blood smear, we use the formula: % Parasitemia = (Number of infected red blood cells / Total red blood cells counted) × 100, with at least ten microscope fields with 500 cells, under an oil immersion microscope [28].

### Outcome Measures and Statistical Analysis

Survival curves were generated and analyzed using GraphPad Prism version 11.0.2. Kaplan-Meier survival analysis was conducted to evaluate differences among groups, with statistical significance determined by Gehan-Breslow-Wilcoxon tests. Parasitemia progression was plotted using GraphPad Prism version 11.0.2 and Unpaired t test with Welch’s correction was conducted to evaluate differences between parasitemia. A p-value <0.05 was considered statistically significant, with further annotations: p < 0.01 (*), p < 0.001 (**), and p < 0.0001 (***).

## Results

Survival analysis revealed significant differences among all experimental groups (p < 0.0001) (Figure 1). In the control group (CTL), 75% of mice died between days 7 and 16 post-infection (Figure 1 and Figure 2), in the absence of neurological signs or pulmonary syndrome. Notably, one male exhibited splenomegaly associated with hepatic involvement and urinary tract occlusion, which led to euthanasia (Figure 3). Microscopic examination of blood smears demonstrated a progressive and continuous increase in parasitemia (Figure 2). The deceased mice exhibited parasitemia levels ranging from approximately 19% on day 7 to around 61% on day 16. The remaining two mice (25%) were euthanized on days 18 and 19 post-infection, respectively, after parasitemia levels had increased more gradually.

**Figure 1:**
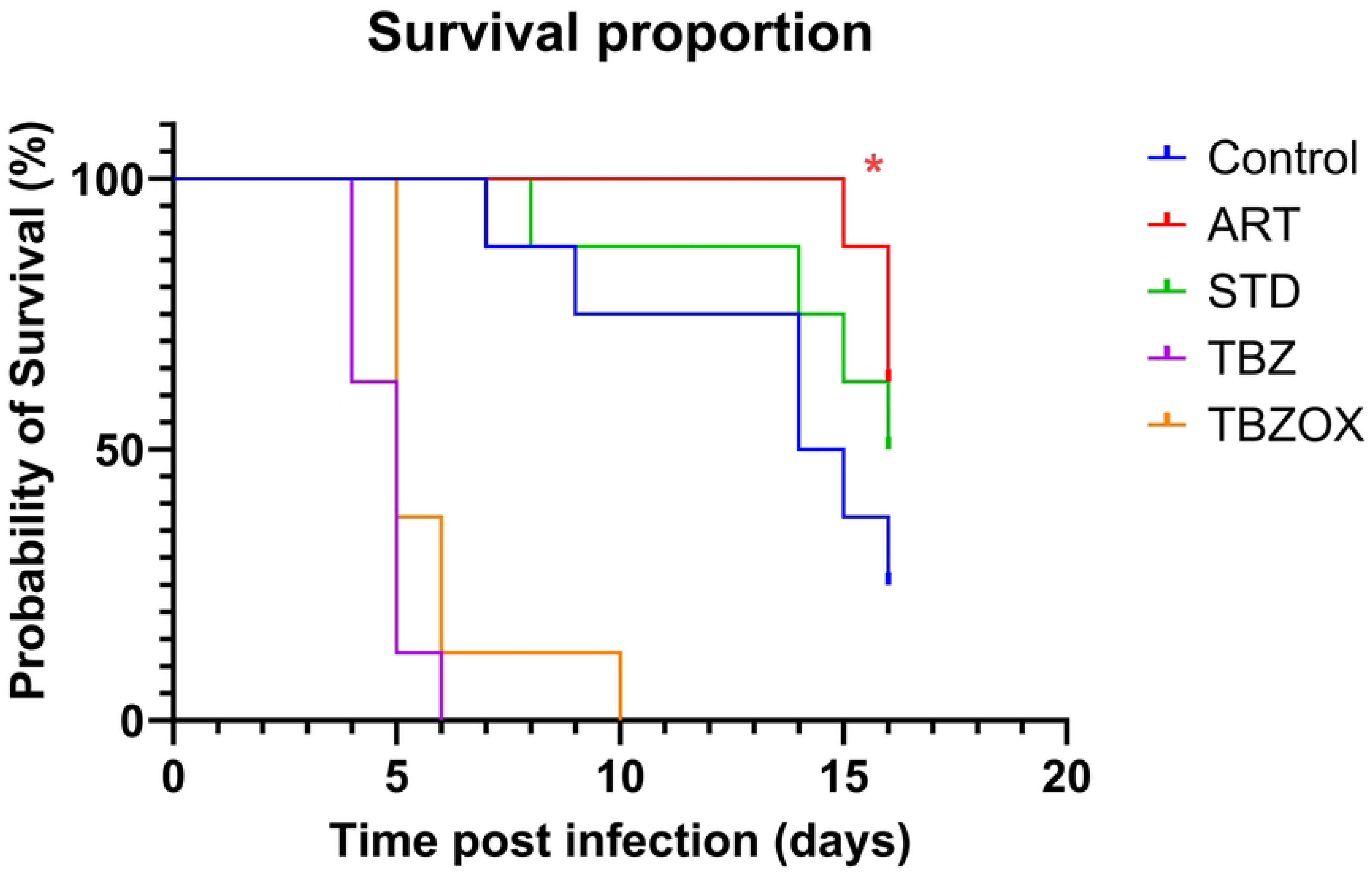
Survival proportion in different groups, from D0 to our experimental endpoint D16. Parasitemia after blood smear is recorded. CTL group is blue, as negative control, ART group is red as positive control, STD group is green, TBZ group is purple and TBZOX group is orange. Only ART group shows efficiency to cure, and it increases survival rate in comparison with CTL group. ART is statistically more effective than other treatments (red asterix).

**Figure 2:**
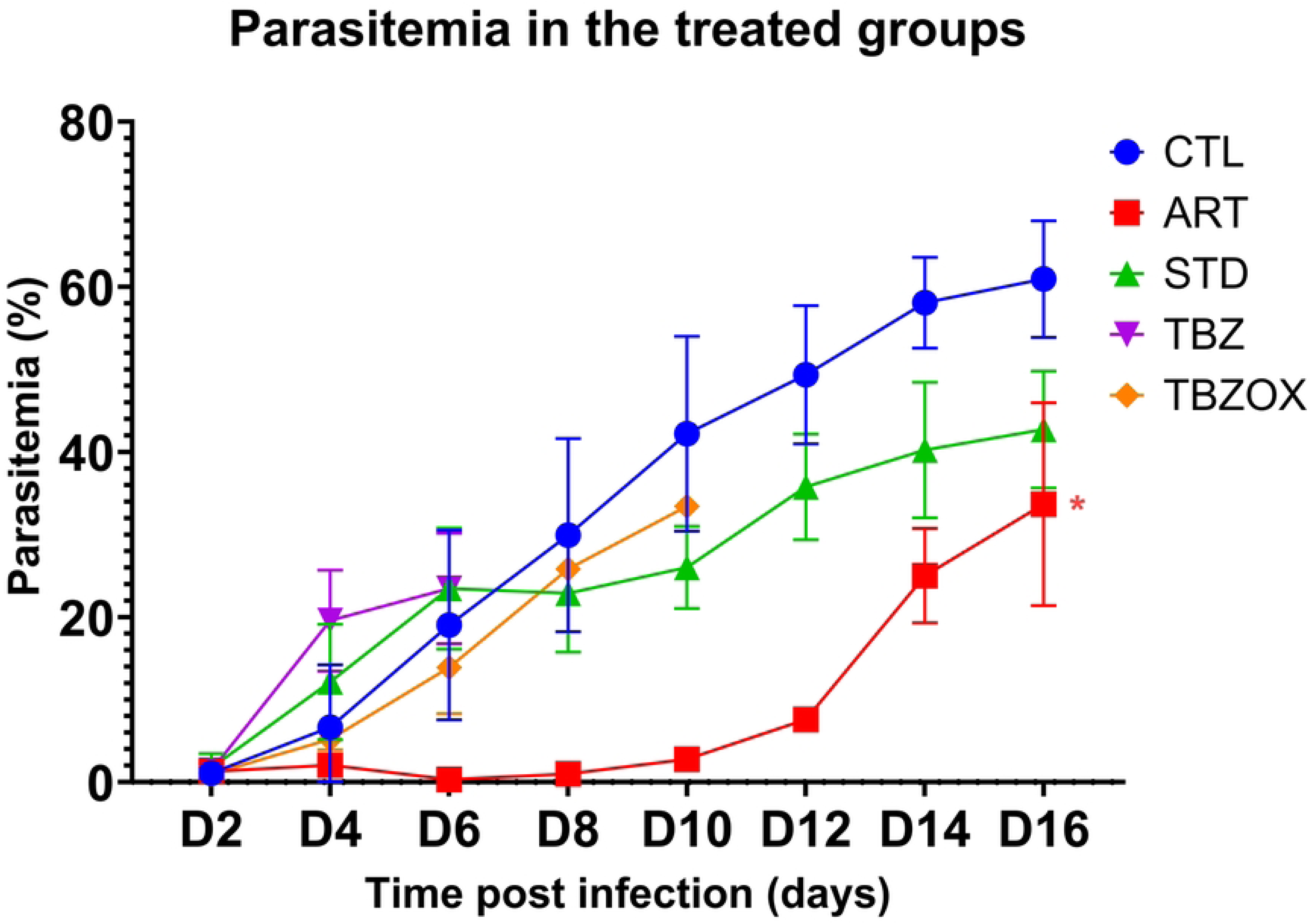
Parasitemia in different groups, from first blood smear at D2 to our experimental endpoint D16. Parasitemia after blood smear is recorded. CTL group is blue, as negative control, ART group is red as positive control, STD group is green, TBZ group is purple and TBZOX group is orange. STD, TBZ and TBZOX are experimental therapies against *Pb*A-infected mice. No experimental drug is under parasitemia from ART group. ART is statistically more effective than other treatments (red asterix).

**Figure 3:**
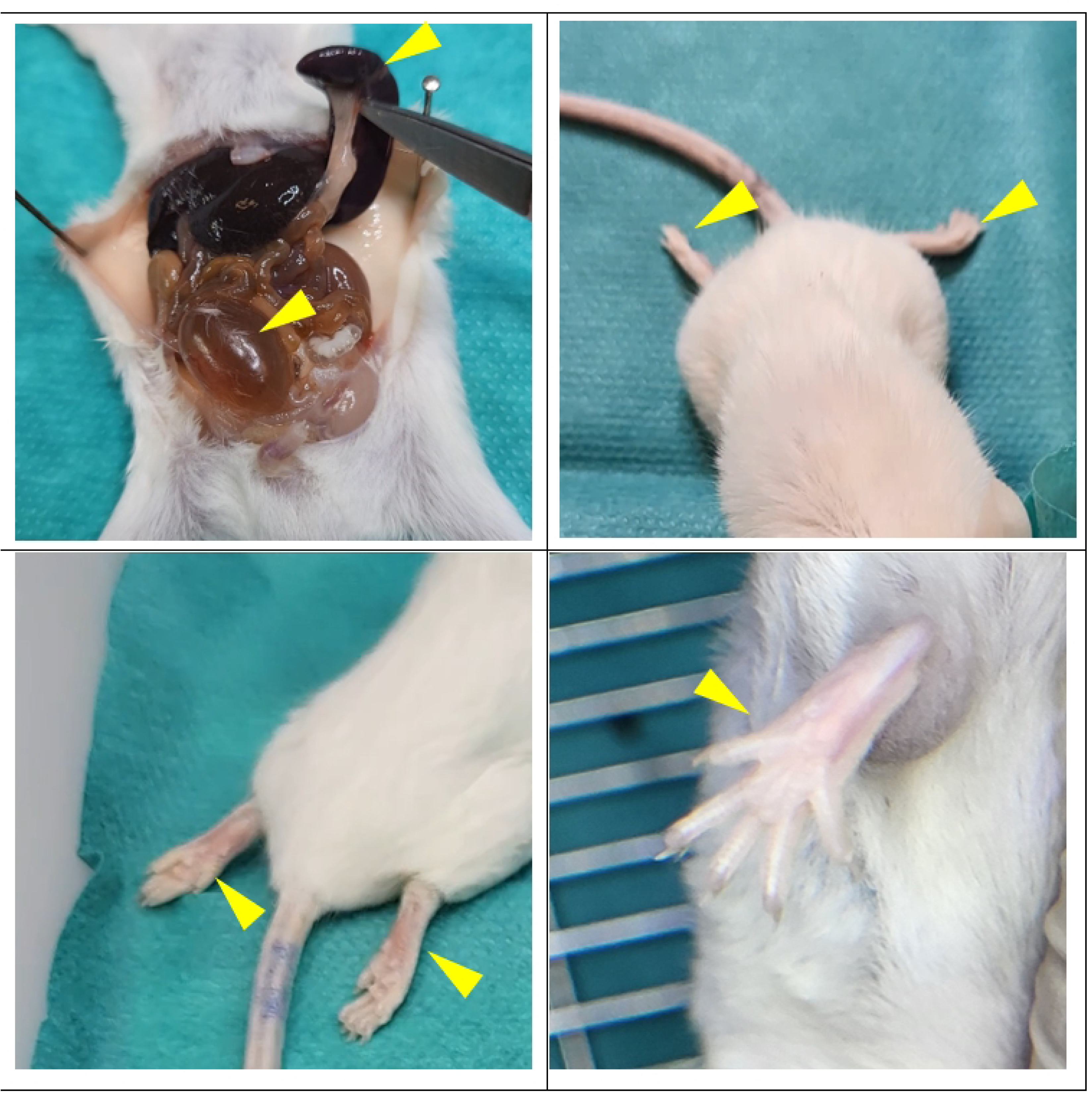
Clinical aspect of mice in the treated group. A splenomegaly in one mouse (male) in CTL group (panel A, top left). Cerebral damage and paralysis in ART group (top right and bottom left). Anemia in STD group (bottom right).

In the artemisinin-treated group (ART), 37.5% of mice succumbed between days 15 and 16 post-infection. The remaining 62.5% of mice, which survived beyond day 16, were euthanized at the end of the experiment due to progressive parasitemia and deterioration of their well-being after day 20. Two cases of note included the occurrence of neuropathology: one male exhibited paralysis of the hind limbs at day 15, and a female displayed both paralysis and dyspnea at day 16 (*Figure 3*). Although rare, neurological involvement has been previously documented in BALB/cJ mice infected with *Pb*A [29], presenting an intermediate clinical profile compared to C57BL/6J mice infected with *Pb*A [30, 31]. Up to day 16, survival in the ART group was significantly different from that in the CTL group (p = 0.0373). Likewise, parasitemia progression in the ART group differed significantly from that in the control group (p = 0.0207). In the STD group, 50% of mice died between days 8 and 16 post-infection (Figure 1). The surviving 50% were euthanized at the end of the study due to increasing parasitemia and associated impact on their health after day 20. Notably, two mice exhibited motor deficits: one male at day 15 and one female at day 16, without neurological signs but with visible anemia (pallor of paws and snout). Up to day 16, survival in the STD group was not significantly different from that in the control group (p = 0.2798). Similarly, parasitemia progression did not differ significantly between STD and control groups (p = 0.4255). The survival duration of mice in the STD group was statistically comparable to that of the ART group (p = 0.3821). However, parasitemia levels in the STD group were significantly higher than those observed in the ART group (p = 0.0284) (Figure 2).

In the TBZ group, all mice died between days 4 and 6 post-infection, without neurological or pulmonary damage. Survival was significantly reduced compared to the control group (p = 0.0002). The parasitemia trajectory in the TBZ group was significantly different from that of the control group up to day 6 (p = 0.0319). Mice treated with TBZ also exhibited significantly shorter survival than those in the ART group (p = 0.0002). Nonetheless, parasitemia levels in the TBZ group did not differ significantly from those in the ART group until day 6 (p = 0.1891). In the TBZOX group, all mice died between days 5 and 10 post-infection, again without neurological or pulmonary signs. Survival analysis indicated a significant difference compared to the control group (p = 0.0005). The parasitemia progression in the TBZOX group was not significantly different from that in the control group (p = 0.6955). Mice treated with TBZOX had significantly shorter survival than those in the ART group (p = 0.0002), and parasitemia levels were notably higher than in the ART group (p = 0.0469).

Overall, none of the three repurposed treatments improved survival or reduced parasitemia levels (Figure 1, and Figure 2). Furthermore, there were no significant differences in efficacy among these treatments (p > 0.05).

## Discussion

This study aimed to evaluate the efficacy of several antiparasitic agents, notably sutezolide (STD), triclabendazole (TBZ), and triclabendazole-oxide (TBZOX), in a murine model of *PbA* infection. Our results indicate that these treatments did not significantly improve survival nor reduce parasitemia, highlighting the challenges in translating antiparasitic effects observed in other models or against different parasites to *Plasmodium* infection.

The initial hypothesis regarding sutezolide’s potential antimalarial activity was based on its documented antimicrobial properties, particularly against *Mycobacterium tuberculosis* [32]. Its mechanism of action, involving inhibition of essential metabolic pathways, could theoretically impact *Plasmodium spp*., which relies on multiple enzymes and biochemical processes. However, to our knowledge, no study has yet confirmed direct antiplasmodial activity of STD, warranting further investigation.

TBZ, a benzimidazole compound, is well-known for inhibiting microtubule polymerization, as demonstrated in *Fasciola hepatica,* where it disrupts cytoskeletal formation and compromises vital parasite functions [33, 34]. Although tubulin sequences differ between Fasciola hepatica and *Plasmodium falciparum,* the structural conservation of tubulins across various protists suggests that microtubule inhibition may represent a valid therapeutic strategy [35]. Indeed, microtubules play critical roles in *Plasmodium* cell division (schizont mitosis) and cytoskeletal formation [36]. *In vitro,* studies have shown that certain benzimidazoles, such as mebendazole, inhibit *Plasmodium* growth [11]. Nonetheless, the lack of direct comparative data on tubulin sequences limits confirmation of this hypothesis, which requires validation through targeted *in vitro* studies [11].

The lack of efficacy observed in our murine model may be explained by multiple factors, including pharmacokinetic properties of the compounds, the parasite’s ability to circumvent cytoskeletal inhibition, or differences in susceptibility between parasites and human targets. Moreover, the dynamics of parasitemia and disease progression in this model differ from in vitro conditions or other experimental systems, complicating result translation. These findings emphasize the need to identify more specific targets and molecules with higher affinity for parasite microtubules.

One of the major challenges encountered in this study was the poor solubility of the tested compounds, which significantly impacted their formulation and subsequent administration. Both STD and TBZ are characterized by limited aqueous solubility, a factor that complicates the preparation of stable and bioavailable formulations suitable for *in vivo* use. Poor solubility can lead to inconsistent dosing, reduced absorption, and ultimately subtherapeutic drug levels at the site of infection. This physicochemical limitation necessitates the use of appropriate solvents or excipients to enhance solubility; however, these additives may introduce toxicity or alter pharmacokinetic profiles, further complicating the interpretation of efficacy results.

Regarding administration routes, although intravenous (IV) and intraperitoneal (IP) injections are commonly employed in experimental models for rapid and controlled drug delivery, they present practical and biological limitations in the context of our study. IV administration requires sterile, water-soluble formulations and precise dosing techniques, which are challenging to achieve with poorly soluble compounds without risking precipitation or embolism. IP injection, while somewhat less demanding in formulation, still involves invasive procedures that can induce stress responses in animals, potentially affecting the immune system and disease progression, thereby confounding the assessment of antiparasitic efficacy.

In contrast, oral administration (*per os*) was chosen for its translational relevance and feasibility despite its own challenges. Oral dosing is non-invasive, reduces animal stress, and better mimics the clinical route of drug administration in humans, which is critical for evaluating pharmacodynamics and pharmacokinetics in a more physiologically relevant manner. Nevertheless, oral delivery of poorly soluble compounds can suffer from low bioavailability due to limited dissolution in the gastrointestinal tract and first-pass metabolism. To partially overcome these obstacles, formulation strategies such as using suspensions, solubilizing agents, or lipid-based delivery systems were employed to enhance oral absorption. However, the inherent limitations of the compounds’ solubility and stability, coupled with the complexity of the gastrointestinal environment, likely contributed to the suboptimal therapeutic outcomes observed.

Overall, the solubility and administration challenges underscore the necessity for developing improved formulations and delivery methods tailored to the physicochemical properties of antiparasitic agents. Addressing these issues is essential to accurately evaluate the true potential of these compounds in preclinical models and to facilitate their translation into effective antimalarial therapies.

Initially based on the literature review by Fonthina *et al*.[37] her propose TBZ for the liver stage, we chose to examine its activity against the blood stage due to the pathophysiology of malaria and the limited range of available treatments in the face of growing resistance to antimalarials. In conclusion, neither sutezolid nor triclabendazole demonstrated significant antiparasitic activity in our *Pb*A – infected murine model at the erythrocyte stage. Nevertheless, microtubule inhibition remains a promising approach given the parasite’s critical dependence on these structures during its cell cycle. Further *in vitro* studies specifically targeting *Plasmodium* with microtubule inhibitors are essential to validate this strategy. Moreover, we fully acknowledge that these results do not close the question of triclabendazole repositioning for antimalarial purposes. In particular, liver-stage activity remains to be investigated *in vitro,* for instance using *P. falciparum* sporozoites infecting HC-04 hepatocyte cultures [38], before a definitive conclusion can be drawn regarding the therapeutic potential of these compounds against *Plasmodium* infection. The development of molecules capable of selectively targeting parasite microtubules may open new avenues for designing effective antimalarial agents.

## List of abbreviations

ART: Artesunate
CTL: control group
IP: intraperitoneal
IV: intravenous
LLINs: long-lasting insecticidal nets
NR3C: National Centre for the Replacement, Refinement and Reduction of Animals in Research
*Pb*A: *Plasmodium berghei* ANKA
STZ: sutezolide
TBZ: triclabendazole
TBZONE: triclabendazole sulfone
TBZOX: triclabendazole sulfoxide
WHO: World Health Organization

## Declarations Ethics declaration

All anesthesia procedures were performed under ketamine and xylazine mixed solution, and all efforts were made to minimize suffering. Euthanasia was performed under anesthesia via cervical dislocation. All procedures were approved by the Animal Welfare Advisory Committee of IHU Méditerranée Infection and the Animal Research Ethics National Committee (C2EA-14), registration number 202203181217615 and in accordance with national guidelines for laboratory animal care.

## Consent for publication

I have the right to post this manuscript and confirm that all authors have assented to posting of the manuscript and inclusion as authors and I confirm all relevant ethical guidelines have been followed,and any necessary Institutional Review Board (IRB) and/or ethics committee approvals have been obtained. This study does not describe the use of any human data, samples, or any research involving human subjects

## Availability of data and materials

All data produced in the present study are available upon reasonable request to the authors

## Authors’ contributions

JD contributed to write the final draft of the manuscript. RA and LM reviewed the final manuscript. JD, RA and LM read blood smears and they analyzed results –JD design the animal testing - JD and RA carried out animal testing-BP secured funding for the research project. All authors read and approved the final manuscript

## Acknowledgements

I would like to address my sincere acknowledgements to Erica Lopez (President of Animal testing Committee, Marseille CE2A14),

## Author’s information

JD is a biomedical researcher who graduated with a doctorate in human pathology and infectious diseases. JD is highly interested in the analysis of mechanisms involved in host-pathogen interactions but also drug therapy to cure infections. He developed a passionate approach to conciliate *in vitro* and pre-clinical studies by using a mouse model for malaria pharmacotherapy assessment.

